# LASSO Based Analysis for Prediction of Prognostic Signature Genes Associated with Breast Cancer

**DOI:** 10.1101/2024.04.02.587421

**Authors:** Souvik Guha

**Affiliations:** Centre for Interdisciplinary Research in Basic Sciences, Jamia Millia Islamia (A Central University), New Delhi, India

**Keywords:** Breast Cancer, Machine Learning, LASSO Regression, Microarray, Prognosis, PPI Network, Signature Genes, Biomarkers

## Abstract

**Background:** Cancer is a genetic disease, where gene alterations play a significant role in the disease onset and pathogenesis. Analysis of the underlying gene interaction pathways could reveal new biomarkers and could also potentially help in the development of targeted drugs for therapeutics. Microarray techniques have emerged as powerful tools capable of simultaneously measuring the expression levels of thousands of genes, making them invaluable in cancer biology research. However, the processing of the resultant datasets poses significant challenges due to their high dimensionality. Also, feature extraction becomes essential to discern the crucial features within these extensive datasets. To mitigate these difficulties advanced computational techniques like Machine Learning (ML) could be instrumental. LASSO-regression-based classification is an advanced ML technique that can help in feature selection by evaluating individual parameters like genes.

**Methods:** This study focuses on uncovering key prognostic genes for breast cancer using a combination of LASSO regression-based classifier and statistical bioinformatics models. Differentially expressed genes (DEGs) were identified using the “Limma” package in R, and significant genes were further filtered using the LASSO-based classifier significance coefficient. Genes common to both methods were considered as the focus of this study. Additionally, Protein-Protein Interaction (PPI) networks of these key genes were constructed using STRING, and hub genes, significant modules, and associated genes were identified using Cytoscape.

**Results:** This study identified CCR8, CXCL11, CCL23, CCL24, CCL28, and CCL21 as signature prognostic genes for breast cancer, revealing a strong association between chemokines and breast cancer pathogenesis. Extensive literature searches were conducted to validate and confirm their prognostic significance in the disease.

**Conclusion:** These findings are pivotal for enhancing our comprehension of the pathways involved in breast cancer. Additionally, they hold promise as novel biomarkers for diagnostic purposes and may also reveal significant therapeutic targets for the management of breast cancer.

## 1. Introduction

Breast cancer is one of the most diagnosed cancers among females globally. In 2020 alone more than 2.26 million breast cancer cases were diagnosed, accounting for 11.7% of total cancer cases. Breast cancer is also one of the major causes of mortality among females worldwide [1]. Breast cancer has surpassed lung cancer to account for 1 in 8 cancer diagnoses and 2.3 million new cases in both sexes [2]. Breast cancer may be classified into five subgroups based on molecular diagnosis namely basal-like, HER2, luminal A, and luminal B [3]. Numerous risk factors contribute to the onset of breast cancer. Gender is often the primary unmodifiable factor since women are more likely than males to acquire breast cancer due to the significant association between specific sex hormones and elevated risk. With over 40% of patients over 65 and over 60% of all breast cancer fatalities occurring in this age range, age is also a key factor in the incidence of breast cancer [5]. A family history of the disease also significantly raises the likelihood of developing breast cancer; around 13–19% of newly diagnosed patients report having a first-degree relative with the disease. On the other hand, modifiable factors like taking certain medications like diethylstilbestrol or undergoing hormone replacement therapy (HRT), significantly raise the probability of developing breast cancer [6]. Studies have confirmed that the consumption of alcohol affects the level of estrogen and thus causes a hormonal imbalance that can elevate the probability of carcinogenesis in females [7]. Like other diseases early diagnosis of breast cancer is very crucial for effective treatment and positive prognosis, with signi:icantly lower mortality rates and higher survival rates observed in patients with smaller tumours at the time of diagnosis [8]. Medical imaging techniques such as Mammography, ultrasonography, magnetic resonance imaging (MRI), scintimammography, single photon emission computed tomography (SPECT), and positron emission tomography (PET) are extensively used for diagnosis [9]. Precision critical care medicine now has more prospects due to recent developments in high-throughput transcriptomic technology, which allow for quick and therapeutically useful gene expression profiling in a matter of hours. In diseases like cancer analysis of the transcriptomic data could reveal intricate molecular pathways and could help in the development of targeted therapeutics. Differential gene expression analysis is a popular computational method for identifying genes whose expressions are significantly different between two phenotypes. This study provides a novel approach for improving the statistical transcriptomic analysis techniques’ outcomes by fusion of Machine learning-based expression profile analysis. In the proposed pipeline [Figure 1] implements a LASSO regression-based identification of the significant genes. The gene expression profiles from the tumour samples belonging to 111 breast cancer patients and 84 control samples. Further studies of the overlapping genes using the PPI network and module analysis five signature genes were identified which demonstrated significant involvement in the disease.

**Figure 1.**
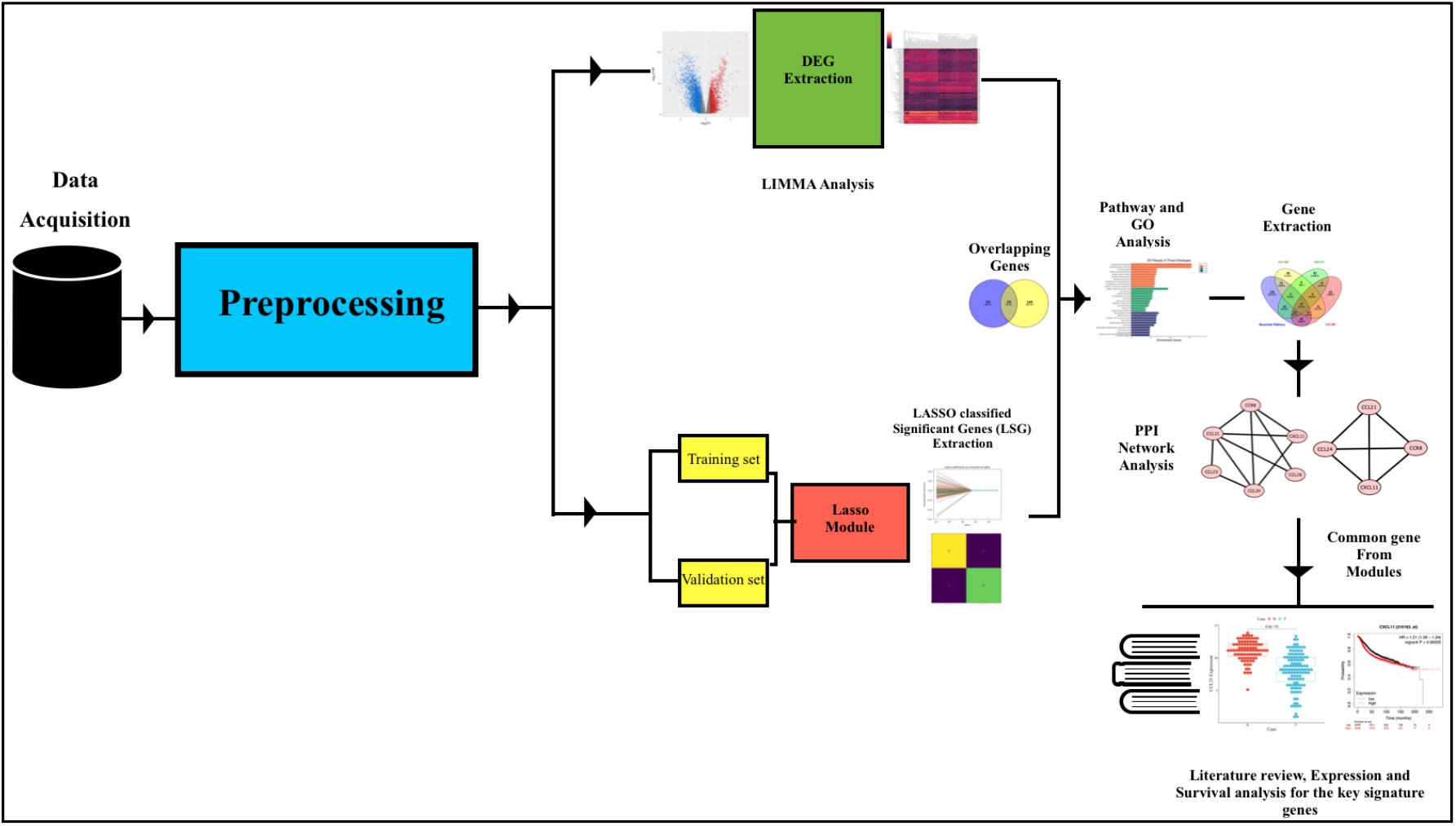
The proposed pipeline of Prognostic Gene analysis from mRNA expression data.

## 2. Material and Methodology

### 2.1 TGCA BRCA mRNA-Seq Data Selection

UCSC Xena browser (http://xenabrowser.net/) was queried for the extraction of The Cancer Genome Atlas (TCGA) breast cancer gene dataset. The data was subjected to back log transformation since the count data were *log*_2_(*x* + 1) transformed. The dataset was then verified with the samples present in the TGCA. Dataset.

### 2.2 Data Validation and pre-processing

Data pre-processing operations included checking for missing values. The edgeR package in R was employed on the mRNA data to obtain the *log*_2_ transformation and the upper quartile(normalization) of the data. The ARSyNseq function of the NOISeq package was applied to this normalized data to perform batch correction. The probe-id mapping with the respective gene symbols was performed using Synergizer [10], gProfile id converter [11], DB2DB conversion [12] and GEO2R [13]. It is typically observed that for genes with several splice variants, many probes match a certain gene symbol. For the purpose of gene mapping, the average expression value of these probes was employed.

### 2.3 Identification of the Differentially Expressed Genes (DEGs)

Following the data preprocessing a *log2* transformation was applied. Then the DEGs were selected using the “limma” [14] package of the R (http://www.r-project.org/). Differentially expressed genes (DEGs) between normal and cancerous samples were defined as those with a fold change greater than two and a p-value less than 0.05.

### 2.4 LASSO-regression Classifier based Significant Genes (LCSG) Extraction

Lasso regression uses the linear regression model as a foundation, but it also applies a technique known as L1 regularization, which adds extra data to the model to avoid overfitting. As a result, we may employ lasso to pick variables by fitting a model with every potential predictor and employing a regularization strategy that reduces the coefficient estimates to zero. Specifically, the reduction target includes the sum of the absolute values of the coefficients in addition to the residual sum of squares (RSS), as in the OLS regression setting.

The residual sum of squares (RSS) is calculated as follows:

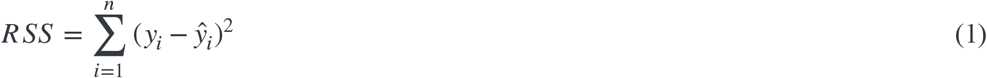

This can also be stated as:

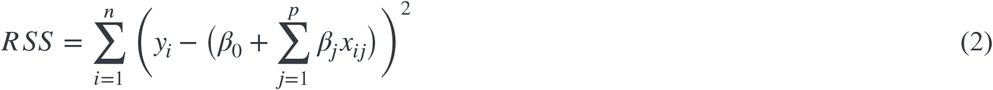

n represents the number of observations, p denotes the number of genes *x*_*ij*_ and represents the value of the jth variable for the ith observation (gene expression level), where i = 1, 2, …, n and j = 1, 2, …, p.

Within the lasso regression, the goal of minimization is transformed into:

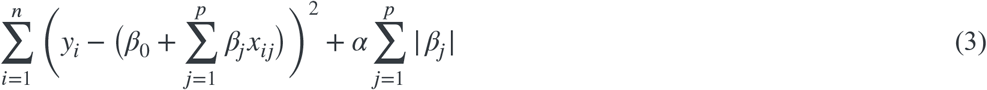

Or

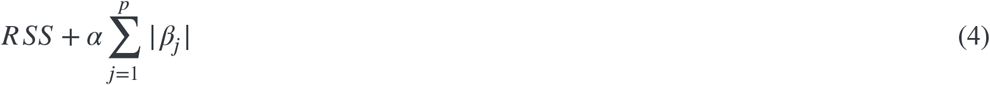

Where α can take various values, depending on the dataset the value is estimated.

The bias-variance trade-off underlies the benefit of Lasso regression over least squares linear regression. The lasso regression fit becomes less flexible as rises, which results in lower variance but higher bias.

We labelled the cancer samples with “1” and the normal samples with “0”. Hence we obtained a binary class dataset. We divided our dataset into a training set and a validation set in a 7:3 ratio respectively using the sklearn train-test split module (https://scikit-learn.org/stable/modules/generated/sklearn.model_selection.train_test_split.html). The procedure for the LASSO-regression-based gene selection is discussed below:

**Step 1:** The optimal α value was estimated using the GridSearchCV module (https://scikit-learn.org/stable/modules/generated/sklearn.model_selection.GridSearchCV.html) and was implemented into the Lasso module of Scikit (https://scikit-learn.org/stable/modules/generated/sklearn.linear_model.Lasso.html).

**Step 2:** The model was trained with the training set.

**Step 3:** The coefficients of all the genes present in the database were calculated.

**Step 4:** The coefficients were sorted in descending order.

**Step 5:** The genes having non-zero coefficients were extracted and other genes were removed.

**Step 6:** The model was evaluated using the validation set and its performance was evaluated, Step 1 was to be repeated if required.

### 2.5 Overlapping Genes Identification between the DEGs and LCSG

To find out the common genes between the differentially expressed genes identified by “limma” and the genes identified as significant by the LASSO classifier, we checked for their overlap, by using the online available tool Venny 2.1.0 (http://bioinfogp.cnb.csic.es/tools/venny/).

### 2.6 Enrichment Analysis of the Overlapping Genes

Based on the GO hierarchy and Reactome 2022 pathway database, we categorized the DEGs into several functional categories of cellular integrals, biological systems, and molecular activities using the online enrichment analysis tool EnrichR (https://maayanlab.cloud/Enrichr/). The top 10 terms of each of these categories were extracted and their overlapping was further studied using Venny 2.1.0 (http://bioinfogp.cnb.csic.es/tools/venny/).

### 2.7 PPI Network Analysis and Significant Module Analysis

The STRING 11.5 (https://string-db.org/) web-based service was employed to create the PPI network on the significant genes [15]. The confidence score was set to greater than 0.70 (very high confidence) and a maximum number of interactions of 0 was set to construct the network [16]. TSV data file was exported from STRING into Cytoscape to construct the PPI network. The proteins are represented by nodes, and protein-protein interactions are represented by edges. A higher node shape signifies a greater degree of connection. The significant modules were identified using MCODE with the default settings: node score=0.2, degree=2, K-score=2 and max depth=100, respectively. The genes in these modules are the signature prognostic genes.

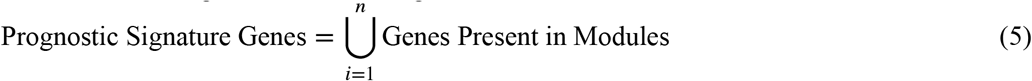

Where n is the number of modules obtained.

### 2.8 Survival Analysis through Kaplan–Meier (KM) plot

The Kaplan-Meier plotter (www.kmplot.com) was employed to assess the predictive significance of signature prognostic genes. This online database has gene expression profiles and survival data of cancer patients. Based on the median gene expression, samples were categorized into high and low-expression groups for OS and recurrence-free survival (RFS) analysis. The log-rank p-value and hazard ratios (HR) were computed along with the matching 95% confidence interval (CI). Significantity was considered as a value of p < 0.05.

## 3. RESULTS

### 3.1 Study Setup

Python version 3.6.9 in the Google Colab environment was used in all the analyses. The operating system was Apple Sonoma 14.2.1 was used in a hardware setup of MacBook Air 2023 based on the Apple M2 chip as the processor and with 8 GB of RAM.

### 3.2 Data Extraction and Identification of the DEGs

The final selected dataset consisted of a total of 30628 genes and 1001 samples out of which 916 were breast cancer patients and the rest 85 were normal individuals. After all the preprocessing operations and gene index mapping the p-value was calculated for the dataset using “limma”. To reduce the dimensionality of the dataset a filter was applied to the p-value to take only the genes with p-value < 0.05 after the DEGs extraction a total of 3858 genes were found to be up-regulated, 8122 genes remained unaltered, and the remaining 2020 genes were also up-regulated, based on the screening criteria of BH-corrected p-value, less than 0.05, and FC (fold change) of more than 2. Volcano plot highlighting the DEGs is shown in Figure 2(a). The variation in the expression levels between normal and cancer patients can be visualized in Figure 2(b).

**Figure 2.**
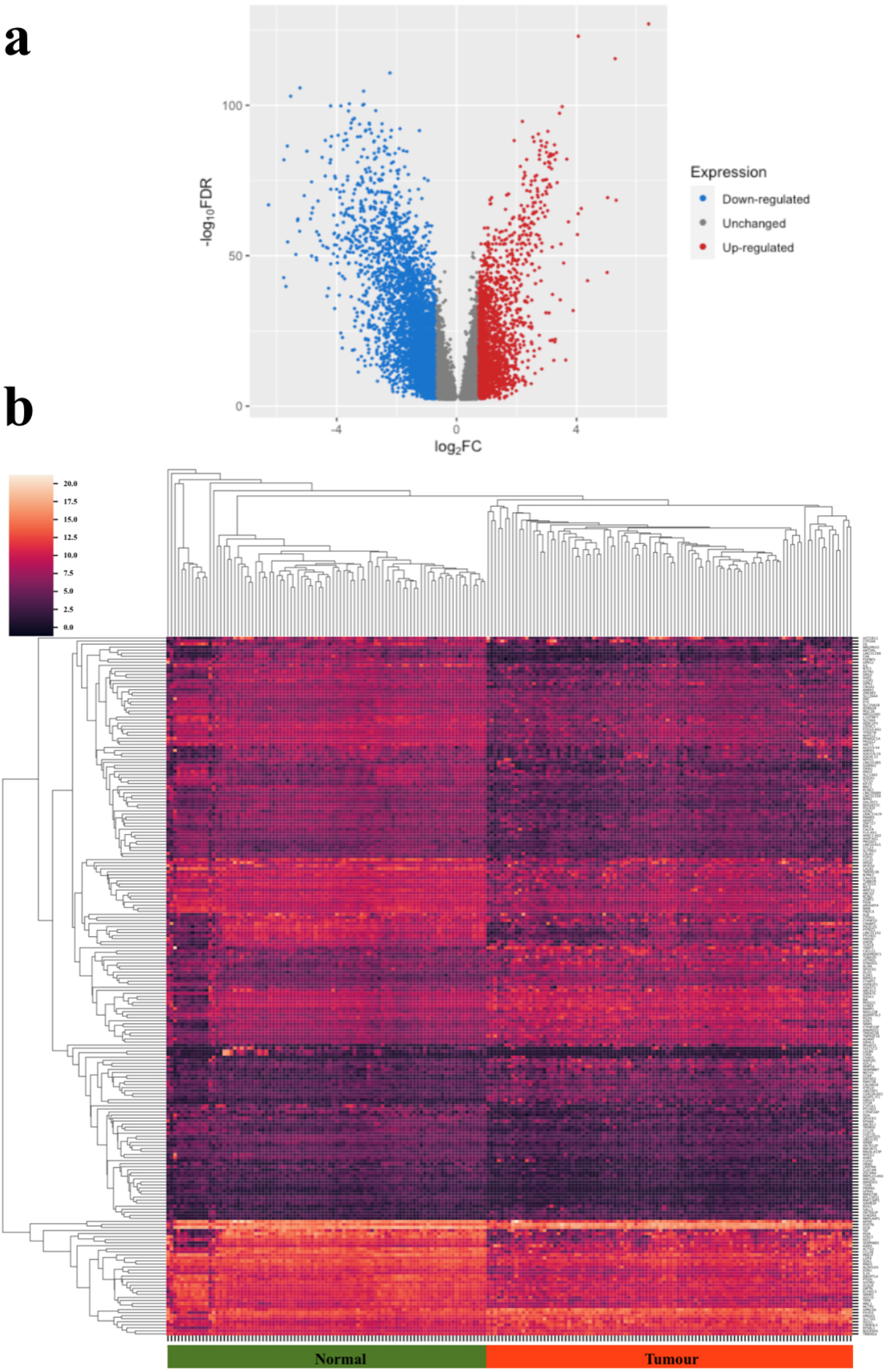
Visualization of the DEGs identified using “limma”. (a) The volcano plot indicates that there are more down-regulated DEGs (shown in blue) than up-regulated DEGs (shown in red), with the unaltered genes (shown in grey). The fold change is shown on the X-axis (log2 scale), and the p-value is shown on the Y-axis (-log10) scale. (b) A heatmap matrix of the DEGs displaying each sample’s expression levels. The sample name is shown on the horizontal axis, while expression level is shown on the vertical axis.

**Figure 3.**
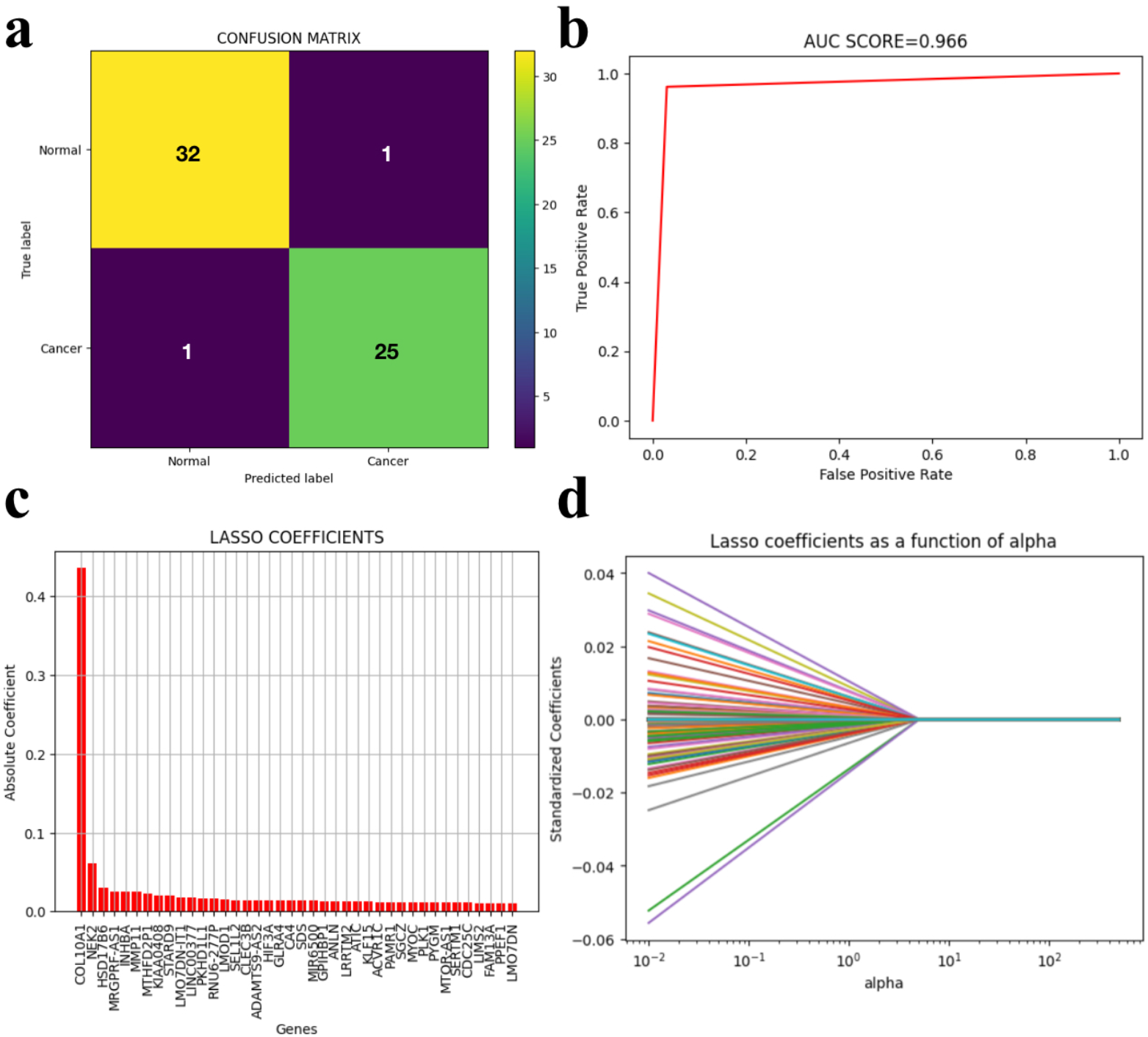
Various evaluation parameters the LASSO-regression classifier. (a) Confusion matrices representation of the results obtained during the validation of the classifier. All the model parameters like accuracy, specificity ROC curves, and AUC value can be calculated using this matrix. (b) The ROC curve of the classifier for different gene subsets(right). This graph’s AUC (Area Under the Curve) score offers an overall performance indicator across all potential classification thresholds. In other words, the AUC score is the probability that the classifier ranks a random positive example higher than a random negative example. (c)Bar plot of the absolute value of the coefficients of the genes in descending order, a higher coefficient signifies a greater weightage during classification. (d) Model coefficients in a regression analysis, with an increase in the penalty parameter (alpha), all weights converge toward zero.

### 3.3 LASSO-regression Classifier based Significant Genes (LCSG)

The LASSO regression classifier was trained with the training set, and the model was evaluated on the validation set. The confusion matrix of the model and the ROC curve were built using the sklearn Confusion Matrix package. The accuracy of the model was evaluated using the accuracy_score module of the sklearn l ibrary (https://scikit-learn.org/stable/modules/generated/sklearn.metrics.accuracy_score.html). Accuracy of a model is defined as :

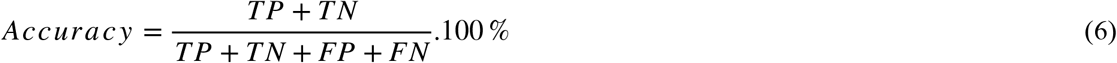

where TP and TN represent true positive and true negative cases while FP and FN represent false positive and false negative cases.

The maximum accuracy of the model was 96.61% and an AUC score of 0.966 for value of 1e-5. Hence value was fixed at 1e-5 for which the classification accuracy was maximum. The coefficients of the genes were extracted using the “coef_” module of sklearn (https://scikit-learn.org/stable/modules/generated/sklearn.linear_model.LinearRegression.html). The absolute value of the coefficients was sorted and all the genes with zero coefficients were removed. Out of the total 140000 genes, 1182 genes had an absolute coefficient value not equal to 0. These 1182 genes were termed LCSG.

### 3.4 Overlapping Genes between DEGs and LCSG

As observed in Figure 4 total of 952 genes were exclusively included in LASSO CLASSIFIED and 1499 genes were included exclusively in the DEGs set identified by “limma”. 230 genes were found to be overlapping between the two sets. These common 230 genes were characterized based on their biological function.

**Figure 4.**
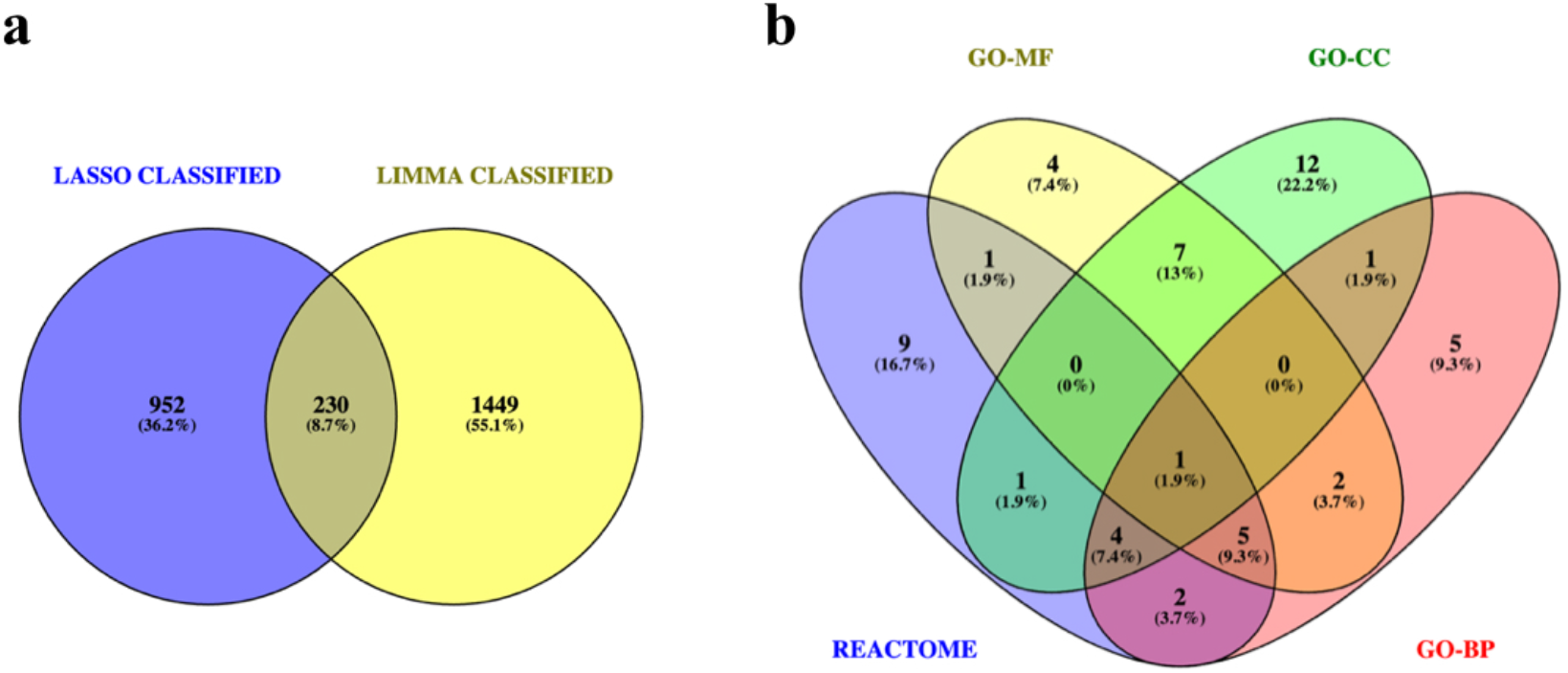
Overlap analysis between various obtained genes sets. (a) Venn diagram representationillustrating the overlap between the gene count classified to be significant by the LASSO Classifier and the genes identified as DEGs using “limma”. As 952 genes were exclusively placed in “LASSO CLASSIFIED”, 1499 genes were exclusively placed in “LIMMA CLASSIFIED”, and 230 genes were mutual to both . Blue circle illustrates the gene count of DEGs in “LASSO CLASSIFIED” group and yellow circle illustrates the gene count in “LIMMA CLASSIFIED” group. (b) Venn diagram representation shows the overlap between the genes associated with the top ten terms of GO Molecular Function (MF), Cellular Component (CC), Biological Process (BP) and the Reactome pathway. As shown, 1 gene was common in all the four sets. The four coloured ovals represent the different sets. Blue represents the genes belonging to the Reactome pathways, yellow represents GO MF genes, green denotes the GO CC genes and red belongs to GO BP genes.

**Figure 5.**
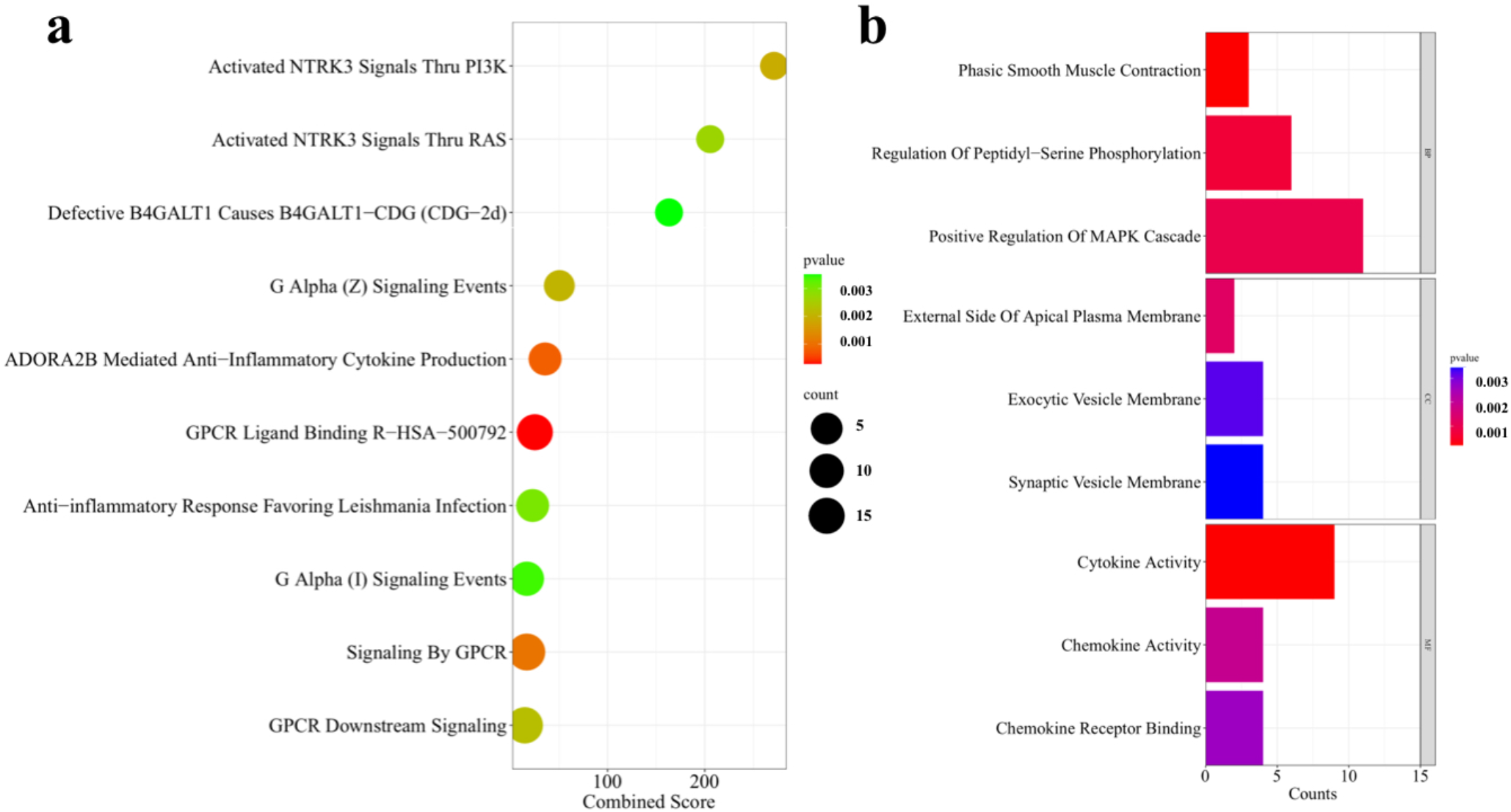
Characterization of the overlapping genes from “limma” and LASSO classifier. (a) Pathways of the genes of interest using Reactome. In EnrichR the Conbined Scores of the different pathways are obtained by multiplying the z-score of the deviation from the predicted rank with the the log of the p-value from the Fisher exact test . (b) GO analysis of the overlapping genes from top Biological Processes(BP), Cellular Components(CC), and Molecular Functions.Counts represent the number of genes present in each category.

**Figure 6.**
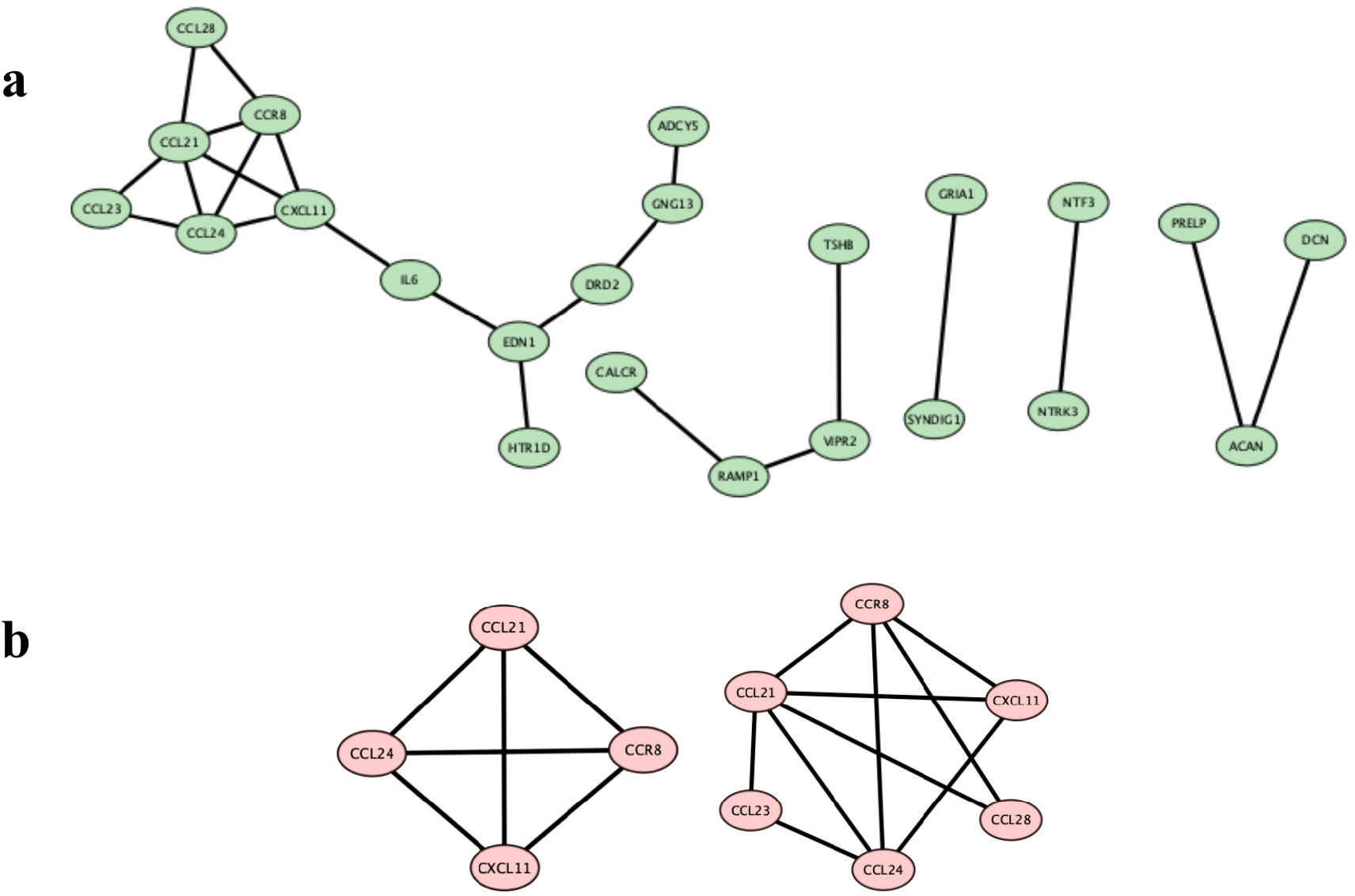
Protein-Protein interaction (PPI) network. This PPI was constructed with the genes present in the top 10 terms of the Reactome, GO-MF, CC and BP respectively using the Venn Illustration in Figure 4. The nodes are the proteins and the edges represents interaction. (a) PPI network as obtained from STRING web module visualized in Cytoscape software. (b) The modules obtained from the network in (a) obtained by applying MCODE on the network. The genes pesent in the modules are the signature prognostic genes.

### 3.5 Characterization of the genes of interest

We utilized the GO hierarchy and Reactome 2022 pathway database to categorize the Differentially Expressed Genes (DEGs) into functional categories of cellular integrals, biological systems, and molecular activities. This categorization was performed using the online enrichment analysis tool EnrichR (https://maayanlab.cloud/Enrichr/), which revealed significant enrichment in the following top three GO terms under each following category: Molecular Function category (terms arranged in descending order of their significance in each category): Cytokine Activity (GO:0005125), Chemokine Activity (GO:0008009) and Chemokine Receptor Binding (GO:0042379). In Biological Process category: Phasic Smooth Muscle Contraction (GO:0014821), Regulation Of Peptidyl-Serine Phosphorylation (GO:0033135) and Positive Regulation Of MAPK Cascade (GO:0043410). In the Cellular component category (descending order): External Side Of Apical Plasma Membrane (GO:0098591), Exocytic Vesicle Membrane (GO:0099501) and Synaptic Vesicle Membrane (GO:0030672). The significantly enriched Reactome pathways (in descending order) are GPCR Ligand Binding (R-HSA-500792), ADORA2B Mediated Anti-Inflammatory Cytokine Production (R-HSA-9660821) and Signaling By GPCR (R-HSA-372790).

### 3.6 PPI Network Analysis and Significant Modules

The PPI Network was created from 55 genes extracted from the top ten GO terms in the three categories and the top ten Reactome pathways were constructed in STRING version 12.0. The network was then exported as a TSV file which was imported into Cytoscape version 3.10.1. Module analysis was done using MCODE. Two Modules each having a score of 3 having 3 nodes, 3 edges and were identified and the genes are termed as the prognostic signature genes on which further studies were carried out on their expression. The genes present in the modules were CCL24, CCL21, CCR8, CXCL11, CCL28 and CCL23.

## 4. Discussion

The morbidity and mortality rates of breast cancer have risen drastically over the last few decades, and there is an urgent necessity to tailor an appropriate management and treatment strategy. A sustained decrease in breast cancer death rates could be achieved with an accurate diagnosis of cancer in the early stages. The most crucial step in achieving the best prognosis is to identify the cells that are cancerous in the early stages. Scientists have investigated numerous methods for the diagnosis of breast cancer, including mammography, positron emission tomography, magnetic resonance imaging, computed tomography, ultrasound, and biopsy. These methods are not suitable for young women and have several drawbacks, including cost and time commitment [17]. For the prediction of diseases like cancer, it becomes crucial to identify the genes that are involved in the disease development and progression. In the analysis of such genes, a Machine Learning technique like the Lasso regression method capable of handling large genetic data like the microarray sequences and adequate statistical knowledge capable of interpreting these data are required. The analysis of the microarray gene expression dataset utilized the methodology of Differential expression analysis and an efficient ML approach to find the common set of significant genes. The DEG analysis compares the degree of expression in the diseased and control groups using a variety of statistical techniques, including the t-test of cohorts. On the other hand, the LASSO-regression classifies the data using its statistical machine-learning approach. While this approach works well, it is limited by the presence of a large amount of noise in the gene expression data, the repeatability of the results, and individual differences due to age, gender, genotype, stage of illness, and other variables. This disadvantage can be removed by implementing a statistical meta-analysis of combined different studies which would enable us to find the distinct disease signatures that are typically consistent in several studies. [18]. Another drawback of using the LASSO-based classifier is that it requires a large set of data for initial training purposes. The dataset should also be balanced which means there should not be any large difference between the counts of the two categories. The LASSO-based classifier implemented in this study demonstrated a significant classification accuracy of 96.6%. From the DEGs identification through “limma” we got a total of 5879 differentially expressed genes. The intersection of these two sets obtained from LASSO and LIMMA gave 230 genes of interest which were used in the further analysis. The genes of interest were subjected to Reactome and GO pathway analysis through Enrichr. The genes were enriched in significant GO terms like Cytokine Activity (GO:0005125), Chemokine Activity (GO:0008009), Chemokine Receptor Binding (GO:0042379), Positive Regulation Of MAPK Cascade (GO:0043410), Neutrophil Chemotaxis (GO:0030593). Earlier studies have shown that Cytokines actively take part in processes involved in tumour onset, promotion, angiogenesis, and metastasis in breast cancer [19]. Angiogenesis creates blood vessels that give the necessary nutrients and oxygen for tumour development in solid malignancies. This mechanism appears to be a well-defined characteristic of cancer and has been shown in several studies to play a critical role in the origin of cancer metastasis. This process is highly controlled, most notably by angiogenesis-promoting or angiogenesis-inhibiting cytokines [20]. Breast cancer is largely caused by persistent inflammation, which is the initial step of malignant growth [21]. Research has indicated that persistent inflammation could increase the probability of developing breast cancer. Additionally, inflammatory regulatory factors secreted by breast cancer cells might facilitate the advancement of inflammation, so establishing a vicious cycle whereby inflammatory tumours enhance the evolution of cancer [22]. These inflammations could trigger inflammatory mediators which result in the production of nonspecific proinflammatory cytokines (TNF-α [tumour necrosis factor-α], IL-6, and IFN-α), which can then trigger the development of chemokines to further inflammation [23]. Studies have shown that in the inflammatory microenvironment, the chemokines and chemokine receptors can support tumour development. According to clinical research, there is a strong correlation between the development of tumours and the overexpression of certain chemokines in breast cancer [25]. Central signalling pathways known as MAPK cascades control a broad range of stimulated cellular processes, such as stress response, apoptosis, differentiation, and proliferation. Consequently, dysregulation, or improper functioning of these cascades, plays a role in the onset and development of illnesses like diabetes, cancer, autoimmune disorders, and faulty developmental processes [26]. Human tissues contain three main MAP kinase pathways; however, the one that involves ERK-1 and -2 is more pertinent to breast cancer. The primary regulators of ERK-1 and -2 are peptide growth factors that function through receptors that include tyrosine kinase. A variety of additional ligands can also function non-genomically by activating MAP kinase through heterotrimeric G protein receptors, including progesterone, testosterone, and estradiol. According to recent research, there is often a higher percentage of cells with active MAP kinase in breast tumours [27]. The most prevalent leukocytes in the body, neutrophils, are becoming more well-acknowledged for their potential to actively modulate cancer. In the bloodstream, neutrophils escort circulating tumour cells to promote their survival and stimulate their proliferation and metastasis. Tumour-infiltrating neutrophils (TINs) are the neutrophils that are found in the tumour microenvironment (TME) and interact with cancer cells [28]. High TIN levels have been linked to advanced histologic grade, tumour stage, and the TNBC subtype [29].

In the REACTOME Pathways, some of the significant terms were: GPCR Ligand Binding (R-HSA-500792), Signaling By GPCR (R-HSA-372790), Chemokine Receptors Bind Chemokines (R-HSA-380108). The biggest family of cell-surface receptors, the G protein-coupled receptors (GPCRs), play a role in the initiation and progression of several malignancies, including breast cancer. Due to their abnormal activation and overexpression, GPCRs are commonly linked to several features of cancer, such as angiogenesis, metastasis, tumour development, invasion, migration, and survival [30]. The chemokine activities also come under significance in the pathway analysis, brief discussion on their activities has been previously discussed.

This study found 6 prognostic signature genes namely CCL24, CCL21, CCR8, CXCL11, CCL28 and CCL23 which all belong to the chemokine family. The relative expression profiles are shown in Figure 7. CCR8 which is a cell surface receptor that belongs to Class A of the G protein-coupled receptor (GPCR) family is observed to be upregulated in breast cancer patients. Studies show that it is well CCR8 is essential for recruiting Tregs to the tumour site and creating an immunosuppressive environment that facilitates tumour escape. This recruiting mechanism can be interfered with by inhibiting CCR8, which may enhance anti-tumor immune responses and slow tumour development [31]. According to current studies, anti-CCR8 antibodies can impair CCR8 activity, which lowers the number of Treg cells accumulating in tumours and interferes with their immunosuppressive role [32]. The KM plot analysis[Figure 8] though shows conflicting results. The Overall Survival(OS) analysis although shows that under expression of CCR8 decreases OS. This could be partially attributed to the fact that at the initial stages of the tumour development, the cytokines could show anti-tumour activity [33]. Further studies in this regard are required. CXCL11 is a chemokine superfamily member of the CXC family and has been found to play a significant role in the development of breast cancer [34]. This molecule has been suggested to be the major ligand for CXCR3 and encodes the protein that activated T-cells need to trigger the chemotactic response. Research shows that CXCL11 activates ERK, which in turn promotes the migratory, invasive, and proliferative activities of MDA-MB-231 cells [35]. It is also exhibited by the KM plot that overexpression of CXCL11 decreases OS. From the analysis of the expression profiles [Figure 7], it is evident that the transcription levels of CCL 21/24/23/28 in breast cancer samples were decreased significantly [36]. This study similar to the previous ones proves that low expression of CCL21 is associated with worst OS. CCL21 is linked to enhanced immunogenicity in breast cancer. CCL24 which exhibits a high expression level in several types of cancer, this study found that it is underexpressed in breast cancer with no significant relation with OS. Again previous works have shown that a reduction in the chemokine CCL23 in hepatic tumours is linked to a poor prognosis for HCC patients and may be a strategy used by the tumour cells to avoid the immune system [37]. The current study also confirms these results evident from the expression profile and Survival analysis. Studies have exhibited Oral squamous cell carcinoma cells with detectable RUNX3 expression levels, CCL28 prevented invasion and the epithelial-mesenchymal transition (EMT). Its suppression of EMT was characterized by enhanced E-cadherin expression and reduced nuclear localization of β-catenin. RARβ expression was elevated by CCL28 signalling through CCR10, which also decreased the interaction between RARα and HDAC1 [38]. In the present study, it has been observed that CCL28 is underexpressed, confirming previous studies [39]. Studies have demonstrated the involvement of CCL28 and CCL27 in the immune system’s anticancer response. These chemokines cause anticancer NK cells to infiltrate the tumour, improving the prognosis through a higher expression of these chemokines in the tumour [40]. The results highly support the fact that the chemokines which modulate immune cell trafficking plays a crucial role in breast cancer tumour microenvironment perturbations. They play a significant role in the immune cell infiltration in the tumour progression and could show anti-cancer properties, as well as some pro-cancer characteristics and thus they play an important role in neoplasia.

**Figure 7.**
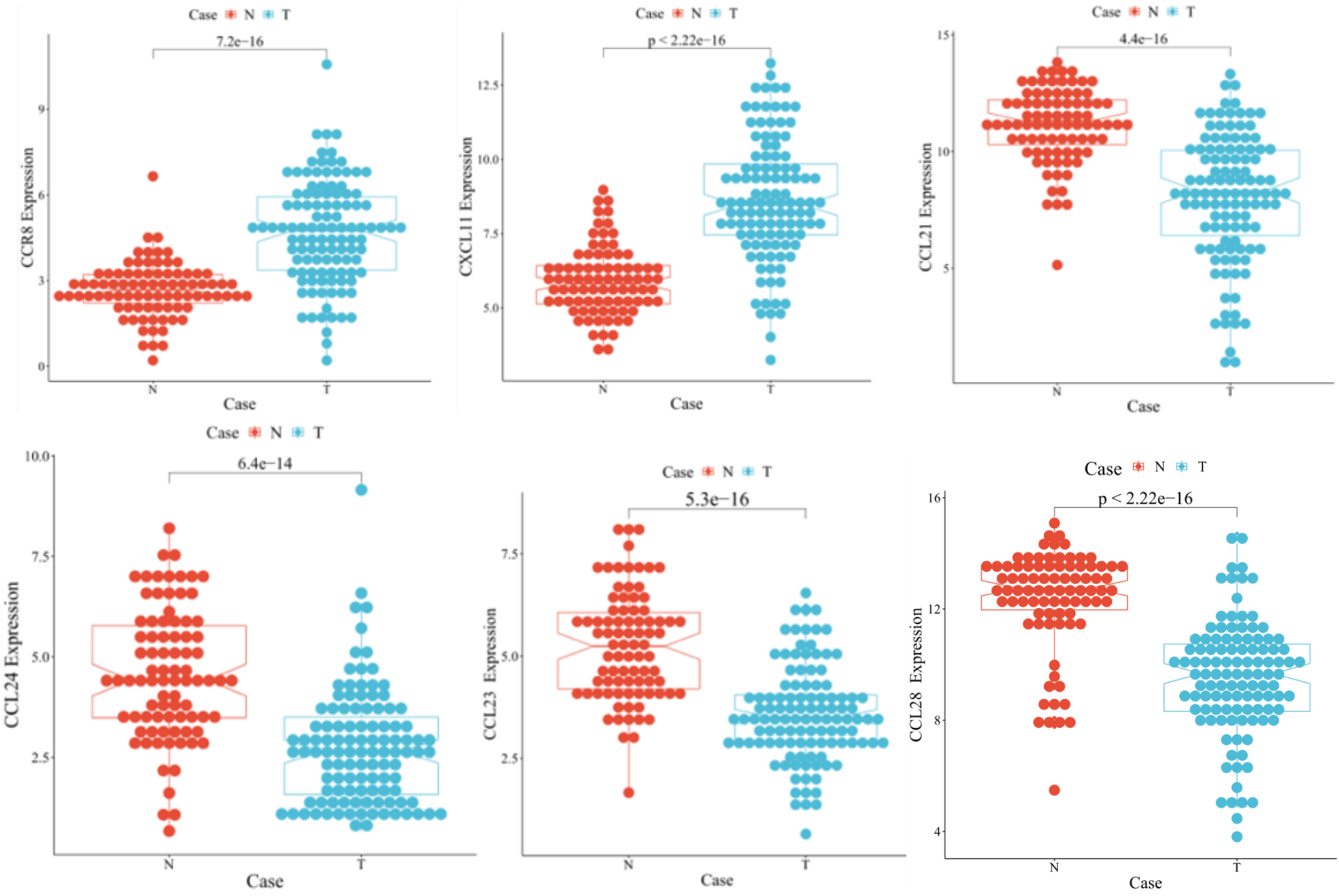
Box Plot depicting Expression levels of the Signature Prognostic genes. The ‘T’ group represents the gene expression in breast cancer patients, while the ‘N’ group represents gene expression in individuals without cancer. The statistical analysis of differential expression between these groups was conducted using the non-parametric Wilcoxon-signed rank test. The expression profile reveals that CCR8 and CXCL11 were notably overexpressed in breast cancer patients, whereas CCL21, CCL24, CCL23, and CCL28 were underexpressed (downregulated) in breast cancer patients. The low p-values obtained from the statistical analysis suggest the potential use of these genes as biomarkers associated with breast cancer.

**Figure 8.**
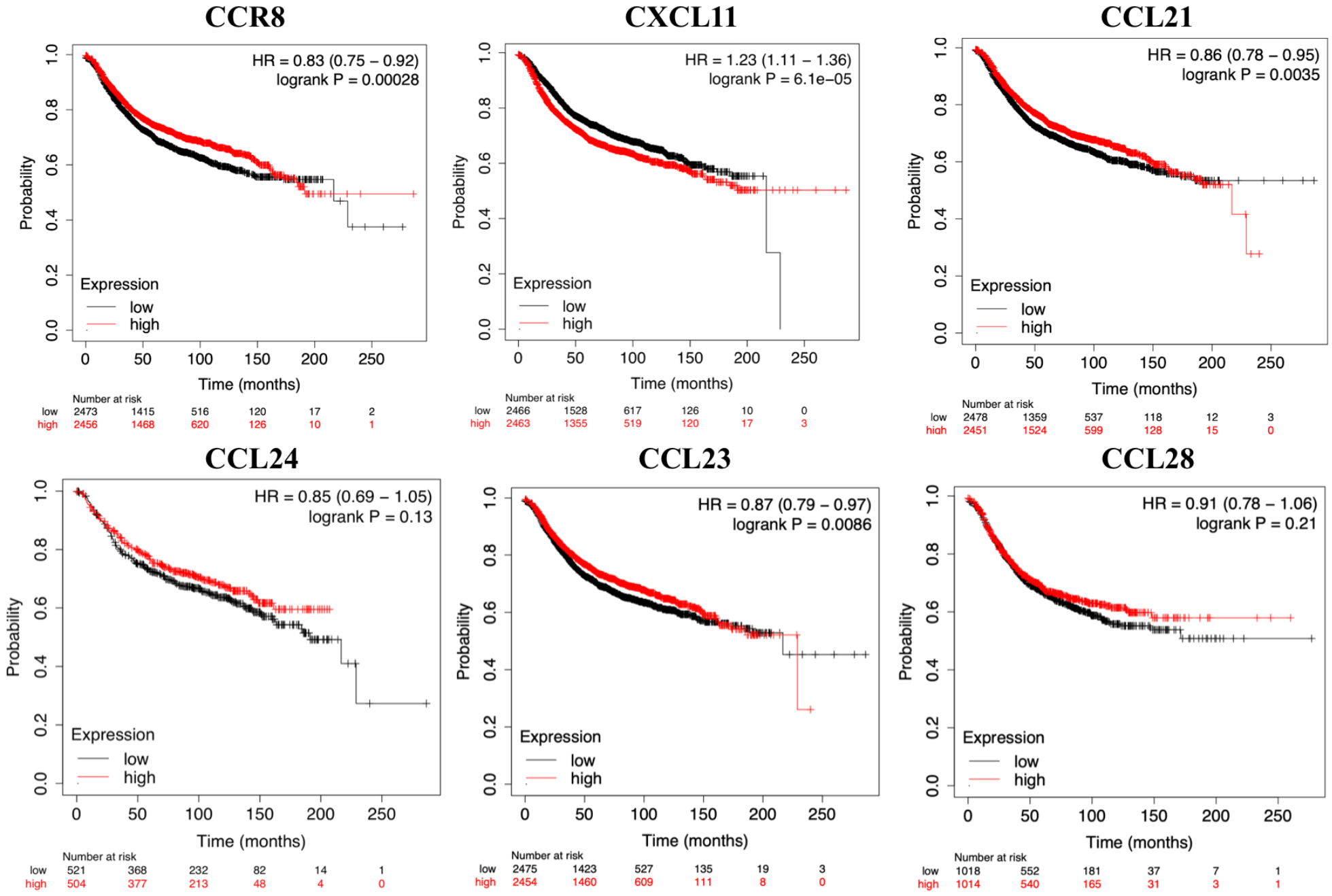
The prognostic value of the CC and CXC chemokines mRNA expression (Kaplan–Meier plotter). In the plots, red and black lines depict the survival curves of patient groups with mRNA expression levels higher and lower than the median levels in the target genes, respectively. The confidence intervals are represented in brackets, and the hazard ratio (HR) is indicated. This analysis provides insights into the potential prognostic significance of the identified signature genes.

## 5. Conclusion

This study has successfully identified several prognostic signature genes and associated mechanisms linked to breast cancer development. While we have pinpointed significant prognostic genes, it’s crucial to recognize that changes in gene expression are governed by regulatory pathways. Understanding these regulatory networks is essential for developing effective disease prediction models. Through the application of various statistical methods and a Machine Learning approach using a Lasso-regression-based classifier, we have unearthed biologically relevant components essential for modelling the intricate associations between genotype and phenotype in breast cancer. These findings will be invaluable to physicians and researchers interested in elucidating correlated pathways and underlying mechanisms in disease development. Further investigations into the roles of these six identified genes in breast cancer development and progression are necessary for enhancing our understanding of the underlying mechanisms.

## 7. Data Availability

The datasets and codes are available in the following GitHub repository: https://github.com/guhasouvik/LASSO_BRCA.git.

## 8. Declarations

### 8.1 Ethics approval and consent to participate

Not Applicable.

### 8.2 Consent for publication

Not Applicable.

